# The *Mitochondrial Iron Regulated* (*MIR*) gene is *Oryza* genus-specific and evolved before the speciation of major AA-genome lineages

**DOI:** 10.1101/846212

**Authors:** Ben-Hur Neves de Oliveira, Andriele Wairich, Andreia Carina Turchetto-Zolet, Janette Palma Fett, Felipe Klein Ricachenevsky

## Abstract

Rice (*Oryza sativa* L.) is both a model species and an economically relevant crop. The *Oryza* genus comprises 25 species, which constitute a genetic reservoir for cultivated rice breeding. Genomic data is available for several *Oryza* species, making it a good model for genetics and evolution within closely related species. The *Mitochondrial Iron Regulated* (*MIR*) gene was previously implicated in *O. sativa* Fe deficiency response, and was considered an orphan gene present only in rice. Here we show that MIR is also found in other *Oryza* species that belong to the AA genome group. We characterized the evolutionary pattern of *MIR* genes within the *Oryza* genus. Our data suggest that *MIR* originated *de novo* from non-coding sequences present only in AA genome species, but these sequences in turn are derived from an exon fragment of *Raffinose Synthase* genes, present in several groups of monocots. We also show that all species that have a putative functional *MIR* conserve their regulation by Fe deficiency, with the exception of *Oryza barthii*. In *O. barthii*, the MIR coding sequence was translocated to a different chromosomal position and separated from its regulatory region, which led to a lack of Fe deficiency responsiveness. Moreover, we show that MIR co-expression subnetwork cluster in *O. sativa* is responsive to Fe deficiency, evidencing the importance of the newly originated gene in Fe uptake. This work establishes that *MIR* is not an orphan gene as previously proposed, but a de novo originated gene within the *Oryza* genus. We also showed that MIR is undergoing genomic changes in at least one species (*O. barthii*), which can impact its role in Fe deficiency.

## Introduction

Asian rice (*Oryza sativa* L.) is one of the most widely cultivated crops, feeding nearly half of the world population and contributing to 20% of dietary calories consumed by humans (Stein, et al. 2018). Rice is also one of the best plant model species, with large resources available for the research community, including reference genome sequences for the two main subspecies, *indica* and *japonica* (Stein, et al. 2018), genome resequencing data for more than 3,000 cultivars and landraces worldwide (Wang, et al. 2018), and mutant databases (Li, et al. 2017). Arguably, rice it is the second most used model plant after *Arabidopsis thaliana*, and based on its synteny with other graminaceous plants, a model for other economically relevant Poaceae species (Gale and Devos 1998).

Rice is a domesticated species, which diverged from its wild progenitors 9,000 to 8,000 years ago (Callaway 2014). The *Oryza* genus includes another domesticated species, *O. glaberrima*, as well as 23 wild relatives, comprising 11 genome types: six diploid (n = 12: AA, BB, CC, EE, FF and GG) and five polyploids (n = 24: BBCC, CCDD, HHJJ, HHKK and KKLL)(Stein, et al. 2018). These species constitute a genetic reservoir which can be searched for interesting genes and alleles to improve useful traits in cultivated rice (Menguer, et al. 2017). To that end, reference genomes from all AA species, which are closely related to rice, and a few distant wild relatives, were recently sequenced (Stein, et al. 2018). These data can also be used to identify orthologous genes of rice in wild species, and thus understand the evolution of sequences in the span of 15 million years comprehending the *Oryza* genus (Zhang, et al. 2019).

Genome sequences from closely related species can also shed light on gene novelty generation within particular lineages. Several genes annotated in any given genome are considered orphans, which means they lack detectable similarity to other sequences in related species (Arendsee, et al. 2014). Characterization of such genes can lead to interesting phenotypes, which may be transferable to other species by simple transgenic insertion (Li, et al. 2009; Li and Wurtele 2015). These unique genes can be generated by different processes, such as extensive divergence from ancestral orthologous genes (Schlotterer 2015), lateral gene transfer (Husnik and McCutcheon 2018), gene loss in related lineages (Zhao, et al. 2015), and *de novo* origination from non-coding sequences (Zhang, et al. 2019). Orphan genes are considered important in the regulation of species-specific traits, including abiotic stresses (Arendsee, et al. 2014). Early work using the rice proteome annotation comprising 59,712 proteins has identified more than 19,000 orphan genes, while a recent study showed that at least 175 genes in Asian rice evolved from non-coding sequences found in wild relatives (Zhang, et al. 2019). These results indicate that newly evolved sequences are quite common in the *Oryza* lineage.

Iron (Fe) uptake in Poaceae species relies on secretion of phytosiderophores that bind Fe^3+^ and are absorbed as Fe^3+^-phytosiderophore complexes, while eudicots and non-Poaceae monocots use Fe^3+^ reduction and Fe^2+^ uptake by specific transporters (Sperotto, et al. 2012). Cultivated rice plants and other Oryza species sharing the AA genome type use a more intricate uptake mechanism, combining features of both strategies (Ricachenevsky and Sperotto 2014; Wairich, et al. 2019). Therefore, this specialization in Fe uptake found in cultivated rice and its close relatives may be aided by newly generated proteins, which can act as modulators of the Fe deficiency response. One of the few rice orphan genes to be functionally characterized to date is named *Mitochondrial Iron-Regulated* Gene (*MIR*) (Ishimaru, et al. 2009). *MIR* was described as having no homologs in other species. The MIR protein was described as being localized in mitochondria. Although its function remains poorly understood, *MIR* was clearly linked to iron (Fe) homeostasis, being transcriptionally upregulated upon Fe deficiency, and repressed under Fe excess (Bashir, et al. 2014; Ishimaru, et al. 2009; Stein, et al. 2019). MIR is regulated by the transcription factor IDEF1 (Iron deficiency element binding factor 1), a master regulator of several of the known Fe deficiency responsive genes in rice roots (Kobayashi and Nishizawa 2012). *MIR* loss-of-function mutants were shorter compared to wild type and were impaired to perceive adequate levels of Fe in tissues, up regulating Fe uptake genes while showing twice as much Fe in roots and shoots (Bashir, et al. 2014; Stein, et al. 2019). Therefore, despite being considered an orphan gene, MIR was shown to be important for Fe homeostasis in cultivated rice plants.

Our work aimed at identifying genes orthologous to *MIR* in closely related wild *Oryza* species and understanding the *MIR* gene evolution, using the available *Oryza* species genome data (Reuscher, et al. 2018; Stein, et al. 2018). Our findings suggest that *MIR* originated before the split of major AA genome lineages, and that *MIR* is not an orphan gene specific to *O. sativa*, but an evolutionary novelty generated within the last one million years in the *Oryza* AA genome lineage.

## Methods

### Data Collection

The omics data (i.e. genomes, proteomes, transcriptomes, and genomic annotation files) used in this work were downloaded for the entire set of available species in both Ensembl Plants (Zerbino, et al. 2018) and Phytozome (Goodstein, et al. 2012) databases. For species occurring in both databases, only the files from Ensembl Plants were kept. This summed up to 89 plant species, from which 10 belong to the *Oryza* genus.

### Retrieving genes homologous to *MIR* by sequence similarity search and phylogenetic analyses

A few search strategies were applied using BLAST (Altschul, et al. 1990). First, the *Oryza sativa* MIR protein sequence (Os12t0282000-01, LOC Os12g18410.1; (Ishimaru, et al. 2009)) was used as a query against the available downloaded plant proteomes using BLASTp and PSI-BLAST strategies (E_value_< 10^−6^) and then against the available downloaded plant genomes using tBLASTn (E_value_< 10^−10^). Afterwards, multiple sequence alignments of both proteins and nucleotide sequences were performed using the MUSCLE software (Edgar 2004).

A conserved portion from the multiple sequence alignment of *MIR* and *MIR* homologous genes was extracted to build a Hidden Markov Model profile (namely, a *MIR-*profile) through the HMMER software (Eddy 1998). The *MIR-*profile was then used to scan the available plant genomes (E_value_< 10^−6^) using the same software. From those hits, sequences longer than 300 nucleotides were selected for phylogenetic analysis. First, a nucleotide multiple sequence alignment was performed using MUSCLE (Edgar 2004) and ambiguous sites were trimmed from the dataset. The jModelTest (Posada 2008) was applied to infer the evolution model that best fit the data and then a Bayesian analysis was carried out using BEAST (Drummond and Rambaut 2007) (“birth-and-death process” was set as a tree prior), running 300 × 10^6^ generations through a Markov chain Monte Carlo process. After that, Tracer (Rambaut, et al. 2018) was used to check for tree convergence. Tree visualization and editing were done using the R packages *Phytools* (LJ 2012) and *ape* (Paradis and Schliep 2019). Finally, domain prediction was performed using the SMART batch perl script provided by the SMART database (Schultz, et al. 2000).

### Mapping IDE elements

For each *MIR* homolog candidate gene, the 2 kb genomic sequence upstream the first CDS codon was retrieved. The occurrence of the IDE1 core element (ATGCT) and its reverse complement was mapped on these genomic sequences using the R package Biostrings (Pagès H 2019). In order to infer statistical significance for the amount of IDE1 elements found in each sequence, a hypergeometric test was applied.

### Synteny and Collinearity Analyses

Synteny analyses among genomic blocks were performed using the McScanX tool (Wang, et al. 2012). A genomic block, in this context, is a chromosomal segment containing 41 genes: *MIR/MIR-like* genes and the 20 first neighboring genes upstream and downstream. First, a BLASTp search is done with the set of proteins encoded by the genes inside the genomic blocks of interest, all against all (E_value_< 10^−6^). Next, the BLASTp output table and the genomic coordinates of the genes in each genomic block are fed to the McScanX tool for synteny inference (E_value_< 10^−10^). Graphical displays of syntenic blocks were made using the R package Circlize (Gu, et al. 2014).

We also performed large-scale multiple sequence alignment (LMSA) with ≈400 kb genomic segments using the LASTz (Harris 2007) aligner followed by MultiPipMaker (Elnitski, et al. 2010), which uses a given reference sequence (RSeq) to guide a multiple sequence alignment. Here, the RSeq used was chromosome 12 of *O. sativa*, from 200 kb upstream to 200 kb downstream of *MIR* (Chr12 - 10451961:10853342). We included in the LMSA all the *Oryza* species available in Ensembl Plants database plus *Leersia perrieri* as an outgroup. The criteria to add a genomic sequence to our LSMA was that it should show at least one of these qualities: i) possesses a putative *MIR* homologous gene; ii) is syntenic to the RSeq. For each species, we found one eligible genomic sequence for the LMSA, except for *O. barthii* and *O. glaberrima* (for which two genomic sequences were eligible). The selection process was carried as follows: we took genomic segment sequences from 200 kb upstream to 200 kb downstream of a given point *P*. If the 200 kb spanning from point *P* went off limits of the chromosome/scaffold, then the expansion was truncated at the border. For the chromosomes with a known or putative *MIR* gene, the point *P* is the center of the *MIR*/*MIR-like* gene. On the other hand, for chromosomes without a known or putative *MIR* gene, the point *P* is the center of maximum potential for synteny with the RSeq. The potential for synteny is here defined as the proportion of tBLASTn hits (E value < 10^-6^), where the queries for this search are the proteins encoded by the set of genes in the RSeq. Therefore, for each target, we looked for the genomic window in which there is the highest density of tBLASTn hits of proteins encoded by the genes that belong to the genomic segment where *MIR* is located. Aligned blocks less than 300 bases and/or less than 50% local alignment identities were discarded.

### Plant Material and Treatments

Rice seeds of *O. sativa* L. (cv. Nipponbare), *O. sativa* ssp. *spontanea* (IRGC 80590), *O. rufipogon* (BRA 00004909-8), *O. longistaminata* (IRGC 101254) and *O. barthii* (IRGC 86524) were submitted to 50 ^◦^C for seven days to break dormancy, according to instructions provided by International Rice Research Institute (IRRI). After, seeds were germinated for four days in petri dishes with two layers of filter paper soaked in distilled water at 28 ^◦^C, two first days in the dark and the two subsequent days in the light (40 μmol.m^2^.s^−1^). After germination, seedlings were transferred to inert soil (vermiculite) for fifteen days. Plants were transferred to plastic pots with 0.5 L of nutrient solution as described (Ricachenevsky, et al. 2011) for seven days, for acclimation. After this period, half of the plants were transferred to the control condition (CC; 100 M Fe^3+^-EDTA) and half of the plants were transferred to the Fe deficiency treatment (-Fe) (no iron added to the nutrient solution). The nutrient solutions were replaced completely twice a week. The pH of nutrient solutions was adjusted to 5.4. Plants were cultivated in a growth room at 24^◦^C ± 2^◦^C under a photoperiod of 16 h day (150 μmol.m^2^.s^−1^)/ 8 h dark. Five days after the onset of Fe deficiency treatment, roots and shoots (n = 4) were collected separately, each sample composed of three plants, frozen immediately in liquid nitrogen and stored at −80^◦^C until processing for RNA extraction.

### RNA extraction and gene expression analysis by RT-qPCR

Total RNA of roots and shoots was extracted using the Plant RNA Reagent (Invitrogen^®^, Carlsbad, CA; USA), following the manufacturer’s instructions, quantified by Nanodrop^®^ (Thermo Fischer Scientific, Waltham, USA) and treated with DNAse I, Amplification Grade (Invitrogen^®^, Carlsbad, CA; USA). cDNA was prepared using OligodT and reverse transcriptase M-MLV (Invitrogen^®^, Carlsbad, CA; USA) following the manufacturer’s instructions, and using 1 µg of RNA. For gene expression analysis, the synthesized first strand cDNA was diluted 100 times. RT-qPCR reactions were carried out in a StepOneTM 7500 real-time cycler (Applied Biosystems, Foster City, California, EUA). One specific primer pair was designed to amplify *MIR* from all species evaluated (F: GCCCATGCTTGCCTTC, Tm value = 59.9^◦^C; R. GCGATATATAGAGGCCACGA, Tm value = 60.1^◦^C; length of amplified fragment: 121 bp). Reactions were conducted as described (Ricachenevsky, et al. 2011). Data were analyzed by the comparative CT method (Livak and Schmittgen 2001). The PCR efficiency from the exponential phase was calculated using the LingReg software (Ramakers, et al. 2003). Ct values were normalized to the Ct value of *OsUBQ5* (F: ACTTCGACCGCCACTACT; Tm value = 61.9 ^◦^ C; R. CTAAGCCTGCTGGTTGTAGAC; Tm value = 61.6 ^◦^ C amplicon size = 62 bp) using the equation as described(Ricachenevsky, et al. 2011). Each data point corresponds to four biological replicates, each replicate composed of roots or shoots from three plants, and three technical replicates.

### Building the Co-expression Subnetwork Around the MIR gene via a Meta-analysis Approach

Two complete sets of microarray experiments done with the Agilent-015241 Rice Gene Expression 4×44K and the Affymetrix Rice Genome microarray technology platforms (both designed for *Oryza sativa*) were downloaded from the Gene Expression Omnibus database (Barrett, et al. 2013). The probe sequences from both platforms were used as queries on a BLASTn (Altschul, et al. 1990) search against *Oryza sativa*’s complete cDNA sequence data in order to annotate both probe sets (applied filters: hit percent identity > 90 and percent coverage > 90). The gene corresponding to the best hit (lowest E-value) of each query sequence was considered to be the target of its matching probe. Exactly one probe was mapped against the *MIR* gene in each platform. The experiments with sample size less than 5 were excluded. This left us with 125 experiments from the Affymetrix platform (total of 3015 microarrays) and 72 experiments from the Agilent platform (total of 2364 microarrays). Data normalization applied to each experiment set of microarrays was dependent on platform technology (RMA for Affymetrix and Quantile Normalization for Agilent).

After that, for each experiment, the following process was carried out: i) the Pearson’s correlation score (PCS) between the *MIR*-matching-probe and every other one was computed; ii) next, we generated a PCS-ordered probe list and selected the top 1,000 ranked probes (*i.e.*, the ones with the 1,000 highest PCS values); iii) for each probe on this group of 1,000 probes, we calculated the Mutual Pearson’s Correlation Score (MPCS) with the *MIR*-matching probe. MPCS is defined as:

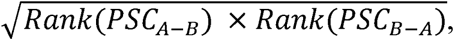

where Rank(PCS_A-B_) is the ranked position of probe B on the PCS-ordered probe list of A; and Rank(PCS_B-A_) is the ranked position of probe A on the PCS-ordered probe list of B; iv) A null distribution of MPCS was drawn through a resampling process, with initial sample size = 1000 and an increment of 500 on each resampling step. This process was repeated until the critical values for statistical significance (p value < 0.01) converged, which here means 10 consecutive resampling steps with less than 10 units difference compared with the previous resampling; v) The pairs of probes for which MPCS values displayed a p value < 0.01 based on the empirical null distribution were considered co-expressed.

The described process generates, for each experiment (*i.e.* each microarray dataset), a set of probes that targets genes that potentially belong to the same co-expressed subnetwork cluster around *MIR*. Next we took the union of these probe sets, filtering out the ones that were not present in at least 5% out of the total number of experiments. Then a single co-expression network was built for each one of the datasets through the *cea* function (number of permutations = 3000; FDR < 0.01) implemented on the R package RedeR (Castro, et al. 2012). The input expression matrix was composed by the resulting union-probeset. The *cea* function outputs an adjacency matrix with weighted edges for statistically significant correlated pairs of gene expression patterns. Negative correlations were set to zero after that. Because every one of these networks have exactly the same set of nodes (the union probeset), we generated a consensus network by averaging non-zero valued weighted edges across all networks. Next, once we verified that the correlation values fitted a normal distribution for the consensus network (Shapiro-Wilk Test, p value > 0.05), we applied yet another filter to keep only the strongest associations of the meta-analysis between probes by zeroing correlations with an associated p value > 0.05. By applying such filtering sequence process, the resulting network degree distributions fitted a power law, which is a known common property of biological networks (Jeong, et al. 2000).

The topological properties of the evaluated graphs and the Gene Ontology term enrichment analyses were computed, respectively, through the R packages igraph (Csardi G 2006) and topGO (Alexa 2019). Node positions in each graph on the 2D grid space (*i.e.* graph layout) were predicted through the Fruchterman-Reingold algorithm (Fruchterman T.M.J. 1991), which tends to maintain close together nodes that share the same neighbors.

In order to investigate how the MIR co-expression subnetwork responds to iron deficiency, we quantified gene expression through a RNAseq analysis with data from Wairich *et al*. (GEO: GSE131238 (Wairich, et al. 2019)). In this study, Wairich *et al*. submitted five weeks old *Oryza sativa* plants to control condition (100 µM Fe EDTA) and iron deficiency (no iron added) for five days. RNAseq library quality control was performed via the FASTQC software. For adaptor sequence removal, we used Trim Galore and the minimum PHRED quality score was set to 30. Filtered reads were aligned against *O. sativa* ssp. *japonica* genome using STAR (Dobin, et al. 2013). Read counting mapped to each gene was performed using the R package GenomicAlignments (Lawrence, et al. 2013). Fold change estimates and differential expression analysis were performed through the R package DESeq2 (Love, et al. 2014).

## Results

### Similarity search identifies *MIR* homologous sequences in the *Oryza* genus

In order to identify potential *MIR* homologs using sequence similarity, we first ran BLASTp and PSI-BLAST using the *O. sativa* MIR protein sequence (Os12t0282000-01) as query against the whole plant proteome datasets from both Ensembl Plants and Phytozome. Both strategies yielded the same hits: ORUFI12G09590 (*OrufMIR*), ONIVA12G08390 (*OnivMIR*), and OBART10G03520 (*ObarMIR*), from *O. rufipogon*, *O. nivara* and *O. barthii,* respectively. Next, we used tBLASTn to find unannotated genomic sequences similar to *OsatMIR*, which expanded our set of *MIR-like* putative genes with the following species: *Oryza longistaminata*, *Oryza glaberrima* and *Oryza sativa* ssp. *indica* (Table 1). With these strategies, we were able to find new putative *MIR* genes exclusively in AA genomes. Most hits were mapped to chromosome 12, with the exception of *ObarMIR*, which is localized in chromosome 10. Unfortunately, it was not possible to obtain chromosome localization for two *MIR-like* hits (namely *OglabMIR* and *OlongMIR*), since those are in unplaced scaffolds (Table 1).

**Table 1.** Putative *MIR* genes found in similarity searches.

| Species | Chromosome/<br>Scaffold | Start | End | Strand | Gene ID | CDS | Protein | Referred<br>as |
| --- | --- | --- | --- | --- | --- | --- | --- | --- |
| *Oryza sativa* ssp.<br>*japonica* | 12 | 10651961 | 10653342 | + | Os12g0282000 | 492 | 164 | *OsatMIR* |
| *Oryza nivara* | 12 | 8237760 | 8238930 | - | ONIVA12G08390 | 447 | 149 | *OnivMIR* |
| *Oryza rufipogon* | 12 | 8920826 | 8921991 | + | ORUFI12G09590 | 444 | 148 | *OrufMIR* |
| *Oryza barthii* | 10 | 3797773 | 3798174 | + | OBART10G03520 | 399 | 133 | *ObarMIR* |
| *Oryza sativa* ssp.<br>*indica* | 12 | 8460341 | 8460766 | - | n.a. | 426 | 142 | *OindMIR* |
| *Oryza*<br>*longistaminata* | KN540552.1 | 951 | 1376 | - | n.a. | 426 | 142 | *OlonMIR* |
| *Oryza glaberrima* | Oglab12<br>unplaced142 | 60166 | 60564 | - | n.a. | 399 | 133 | *OglaMIR* |

n.a. = not available

### The *cis*-acting element IDE1 core sequence is enriched upstream *MIR* genes

The presence of a high number of IDE1 core sequences (CATGC) in the *OsatMIR* promoter region has been reported (Ishimaru, et al. 2009), indicating that IDEF1, a transcription factor that regulates Fe deficiency responses (Kobayashi, et al. 2007), binds to the *OsatMIR* promoter to up regulate its transcription under low Fe availability. In order to investigate whether *MIR* homologous genes found in all seven species (Table 1) have the same IDE1 enrichment to be regulated by IDEF1, we counted how many times IDE1 elements occur within 2 kb sequences upstream of the start codon of each putative *MIR* gene. Assuming a hypergeometric distribution, the number of IDE1 elements in these sequences were statistically enriched (p value < 0.01), indicating that *MIR* homologous candidate genes might also be regulated by IDEF1 and therefore up regulated under Fe deficiency. The exception was *ObarMIR* from *O. barthii*, which did not show such enrichment (p > 0.05). Therefore, we searched the *O. barthii* genome using *MIR* and *MIR* homologous promoter sequences (BLASTn; E value < 10^-10^) and found one hit in chromosome 12, which is enriched with IDE1 elements (hypergeometric test, p value < 0.01). These results are summarized in Figure 1, showing a graphical display of the multiple sequence alignment of *MIR* genes plus 2 kb long genomic sequences upstream of their start codons.

**Figure 1.**
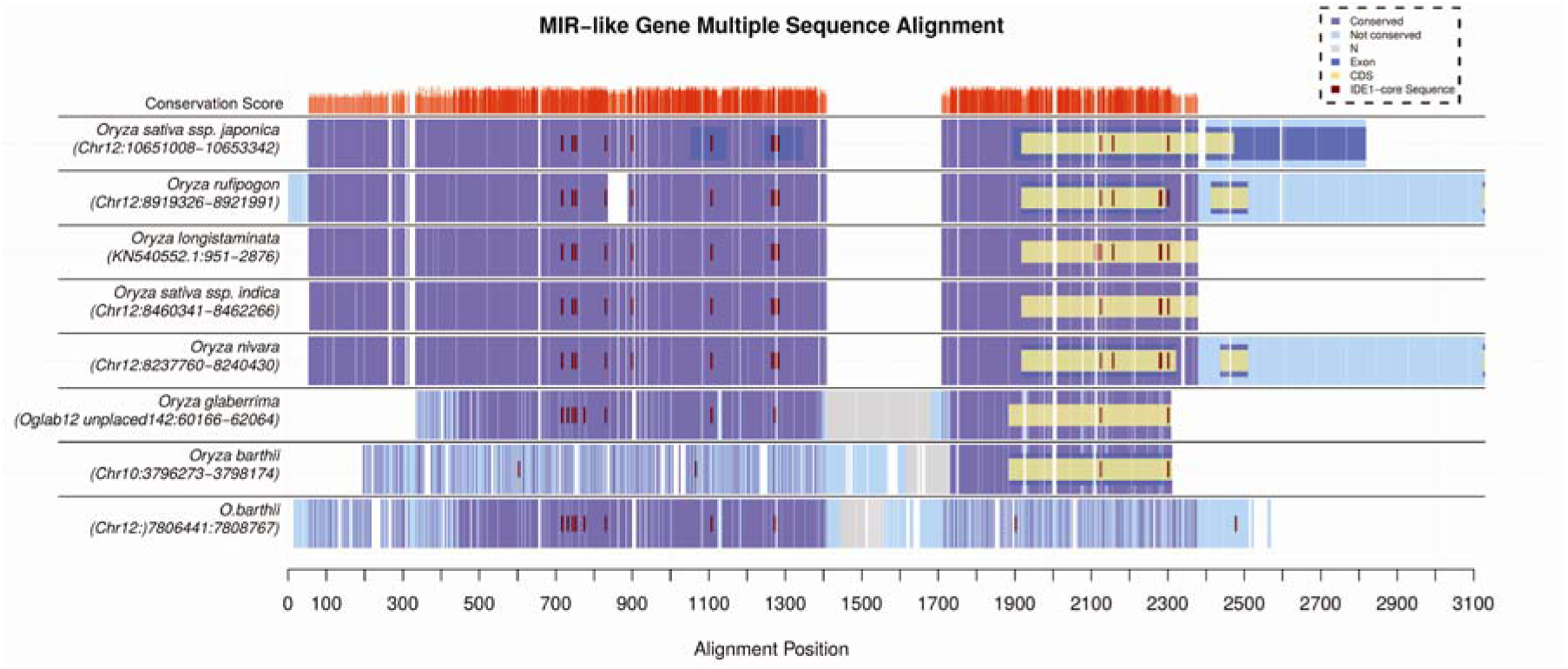
Graphical representation of the genomic DNA multiple sequence alignment of *MIR-like* genes. On the top row, red bars indicate conservation score of the alignment, where conservation score indicates the proportion of nucleotides on that particular site that are equal to the consensus nucleotide for that position. Purple represents conserved sites on the sequences (>50% conservation), while light blue represents non-conserved sites. Annotated exons and CDS (according to ENSEMBL plants annotation files) are shown in blue and yellow, respectively. Undetermined nucleotides (“N”) are grey and blank spaces represent gaps. Short vertical dark red lines indicate where the IDE1 core sequence occurs across the alignment. Sequences from distinct *Oryza* species are identified, and their chromosome and genomic position is shown.

The IDE1 overall distribution is conserved across *Oryza* species that potentially have *MIR* homologous genes (Figure 1). In *OsatMIR*, the first two exons (which are 5’UTRs) and the second exon are especially enriched with IDE1 elements, indicating a possible regulation by IDEF1. The genomic segment comprising these exons is conserved across the studied species, although the homologous genomic sites are not annotated as exons in the predicted gene structures from Ensembl Plants database for *OrufMIR* (ORUFI12G09590), *OnivMIR* (ONIVA12G08390) and *ObarMIR* (OBART10G03520).

While summarizing these results, one would take notice that even though *ObarMIR* is mapped to chromosome 10, there is a genomic fragment on chromosome 12 of *O.barthii* that is highly similar to the IDE1-enriched region of the *MIR-like* genes found in other species, which suggests that while the *ObarMIR* coding sequence is located in chromosome 10, the original promoter region is located in chromosome 12 (Figure 1).

### Synteny in *MIR-like* regions among *Oryza* species chromosomal segments

Synteny analyses revealed that putative *MIR* genes are located in syntenic positions across the *Oryza* species, with the exception of *O. barthii* (Figure 2A). However, the genomic segment in *O. barthii* that holds an identified IDE1-enriched region on chromosome 12 shows high synteny with the genomic segments that hold putative *MIR* genes in the other *Oryza* species (Figure 2A), indicating that the region where the *ObarMIR* promoter is found is syntenic to the regions where *MIR* homologous candidate genes are located. Conversely, it is shown by the same method that the *O. barthii* genomic segment on chromosome 10 carrying the *ObarMIR* gene is syntenic to homologous genomic segments in chromosomes 10 of other *Oryza* species (Figure 2B). These data strongly suggest that *ObarMIR* (OBART10G03520) moved from chromosome 12 to chromosome 10 with minimal disturbance of the genomic environment in its vicinity, while much of the regulatory region of *ObarMIR* remained in chromosome 12.

**Figure 2.**
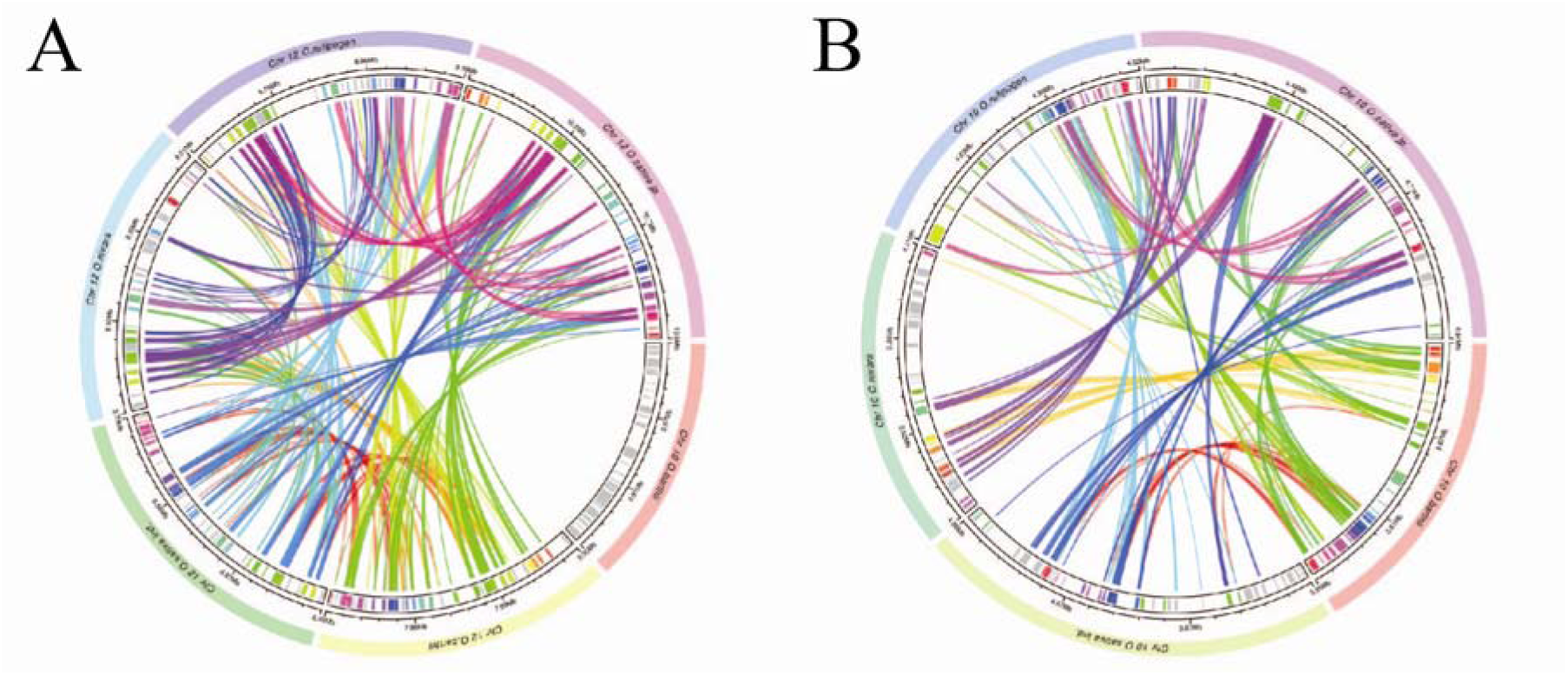
Circular plots of synteny in the *MIR-like* genomic regions. The rectangular arcs represent genomic segments. Vertical bars within arcs represent genes in the genomic segment. Ribbons connecting a pair of genomic loci show syntenic pairs of genes (E-value < 10^-30^). **(A)** Circle plot containing exclusively genomic segments harboring a *MIR-like* gene in *Oryza* species. Homologous genes are represented by vertical lines in the same color (*O. sativa* ssp. *japonica* is the reference, and homologous genes are colored accordingly). Grey lines represent genes with no similarity. **(B)** Circle plot showing the region in chromosome 10 where *O. barthii* has a *MIR* gene. The only arc containing a *MIR-like* gene in this scenario is *O.barthii*’s. Homologous genes are represented by vertical lines in the same color (*O. barthii* is the reference, and homologous genes are colored accordingly). Grey lines represent genes with no similarity.

### Large-scale genomic alignment reveals chromosomal inversions around the ***MIR loci***

In order to further explore collinearity on the genomic context in which *MIR* genes are occurring across *Oryza* species, we performed large-scale DNA alignments of ≈400 kb genomic segments carrying a *MIR* gene and/or being syntenic to a reference chromosome segment (*O. sativa* ssp. *japonica* chromosome 12, from 200 kb upstream to 200 Kb downstream of *OsatMIR*). As shown in Figure 3A, the genomic segment under analysis is fairly conserved in AA genomes, with a few small segments still preserved in more distantly related species such as *O. punctata*, *O. brachyantha* and even *Leersia perrieri*, an outgroup of the *Oryza* genus. The only species that appears to have large-scale collinearity with *O. sativa* ssp. *japonica* is *O. rufipogon*, as essentially the whole genomic block is aligned in the same orientation. We found two major inversions around the *MIR* loci in the other species with AA genomes. The first one is observed in *O. sativa* ssp. *indica* and *O. nivara*, where the inverted segment comprises *MIR* plus the first gene downstream (Os12g0282400). The second inversion is observed in *O. barthii*, *O. glaberrima*, *O. longistaminata* and *O. meridionalis* (the latter does not have a *MIR-like* gene), with the inverted segment comprising one gene downstream and three other genes upstream of *MIR* (Os12g0281600, Os12g0281450, and Os12g0281300) (Figure 3A). In Figure 3B, we confirm that the orientation of the genes inside the inverted genomic blocks is in fact opposite in comparison with our reference chromosomal segment (from *O. sativa* ssp. *japonica*). As already stressed, the most striking case is in *O. barthii*, where the alignment depicts an almost precise removal of *ObarMIR* from chromosome 12 and its insertion on chromosome 10, which comes into agreement with our previous synteny analyses. In *O. longistaminata* and *O. glaberrima*, the genomic segment holding *MIR* is also poorly conserved compared to other species. However, in these two species, the *MIR-like* gene is located at a scaffold and an unplaced contig, respectively. Therefore, it is not possible to assign sure positions of these genes within the genomes of *O. longistaminata* and *O. glaberrima*. It is worth to notice a collinear block downstream *OlongMIR* on the genomic alignment (Figure 3A), which could indicate that this scaffold may belong to chromosome 12 in this species. Nevertheless, whole genome assemblies for these species must be improved in order to allow higher resolution of such syntenic relationships.

**Figure 3.**
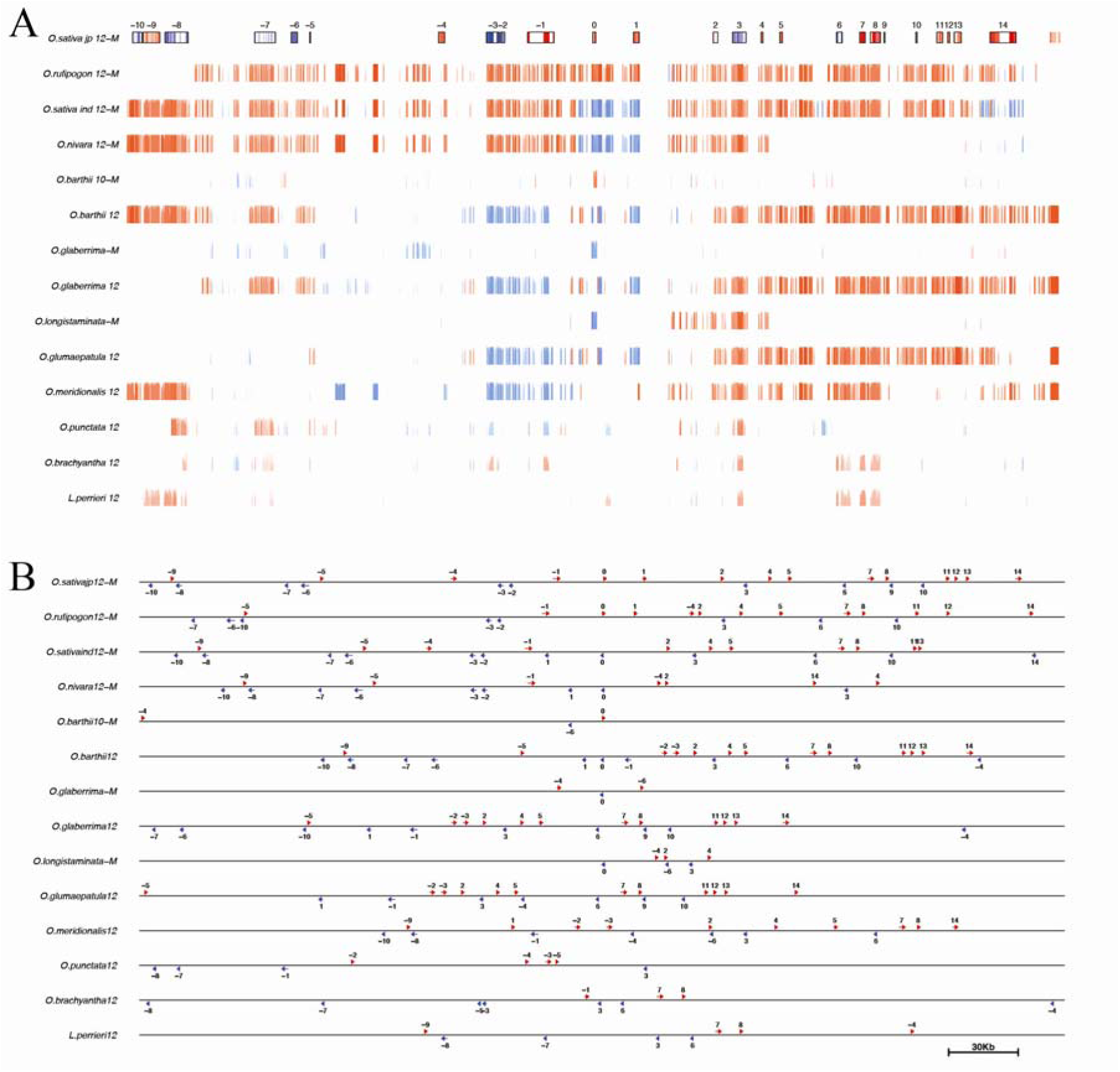
Large Genomic Segments Multiple Sequence Alignment. **(A)** Graphical representation generated using the MultiPipMaker alignment tool. Boxes on the top row represent genes of the sequence used as reference to guide the alignment (*O. sativa* ssp. *japonica* as a reference). The red and blue colors indicate gene orientation (red for forward; blue for reverse); blank spaces represent exons. The numbers above each box give their relative position (negative numbers are upstream the *OsatMIR* while positive numbers are downstream). The red/blue bars below each genomic site represent an alignment of a block containing at least 300 nucleotides and sequence identity > 50% in a local alignment (bar height is proportional to the alignment percent sequence identity). Genomic segments from different *Oryza* species are identified, and segments labeled with “-M” have a *MIR-like* sequence. **(B)** Genomic segments in A, not aligned. The arrows indicate BLASTn hits (E value < 10^-10^) in the sense (red) or antisense (blue) orientation in reference to *O. sativa jp.* The queries are the genes comprised on the *O. sativa* ssp. *japonica* genomic segment and the best hits on the other ones are labeled according to the query on the top row. Black rectangles indicate loci where there are inverted sequences in reference to *O. sativa* ssp *japonica*.

### Fe deficiency up regulates the expression of *MIR* genes in species of the *Oryza* genus

Gene expression analysis has revealed that transcripts of *OsatMIR* in roots from plants grown under Fe deficiency were more abundant compared to shoots and roots grown under Fe sufficient conditions (Ishimaru, et al. 2009). We also observed that IDE1 enrichment is common in *MIR* genes in *Oryza* species (Figure 1). To understand the role of *MIR* genes we evaluated the expression level of these genes in plants of *O. sativa* ssp. *japonica*, *O. rufipogon*, *O.barthii* and *O. longistaminata*. Plants were cultivated as described in Material and Methods and submitted to Fe deficiency for five days. Expression levels of *MIR* genes in roots of *O. sativa* ssp. *japonica*, *O. rufipogon* and *O. longistaminata* were higher when plants were grown under Fe deficiency (-Fe) than in control conditions (Figure 4). The transcript levels of *OsIRT1*, which is involved in Fe^2+^ uptake in *O. sativa* plants (Eide, et al. 1996), *OsNRAMP1*, which is known to be induced in *O. sativa* by low Fe conditions, and YSL15, a Fe^3+^-deoxymugineic acid transporter of *O. sativa* (Inoue, et al. 2009; Lee, et al. 2009), were used as positive controls, indicative of true low iron status of the plants. We observed an increase in transcript levels for all these genes in all species, with the exception of *IRT1* in roots of *O. longistaminata*, which was not upregulated (Figure 4). In the species *O. barthii*, there was no difference in *ObarMIR* expression in roots grown under Fe sufficiency and Fe deficiency conditions. These results indicate that *MIR* genes up regulation under Fe deficiency is conserved, with the exception of *ObarMIR*, which lacks a regulatory region enriched in IDE1.

**Figure 4:**
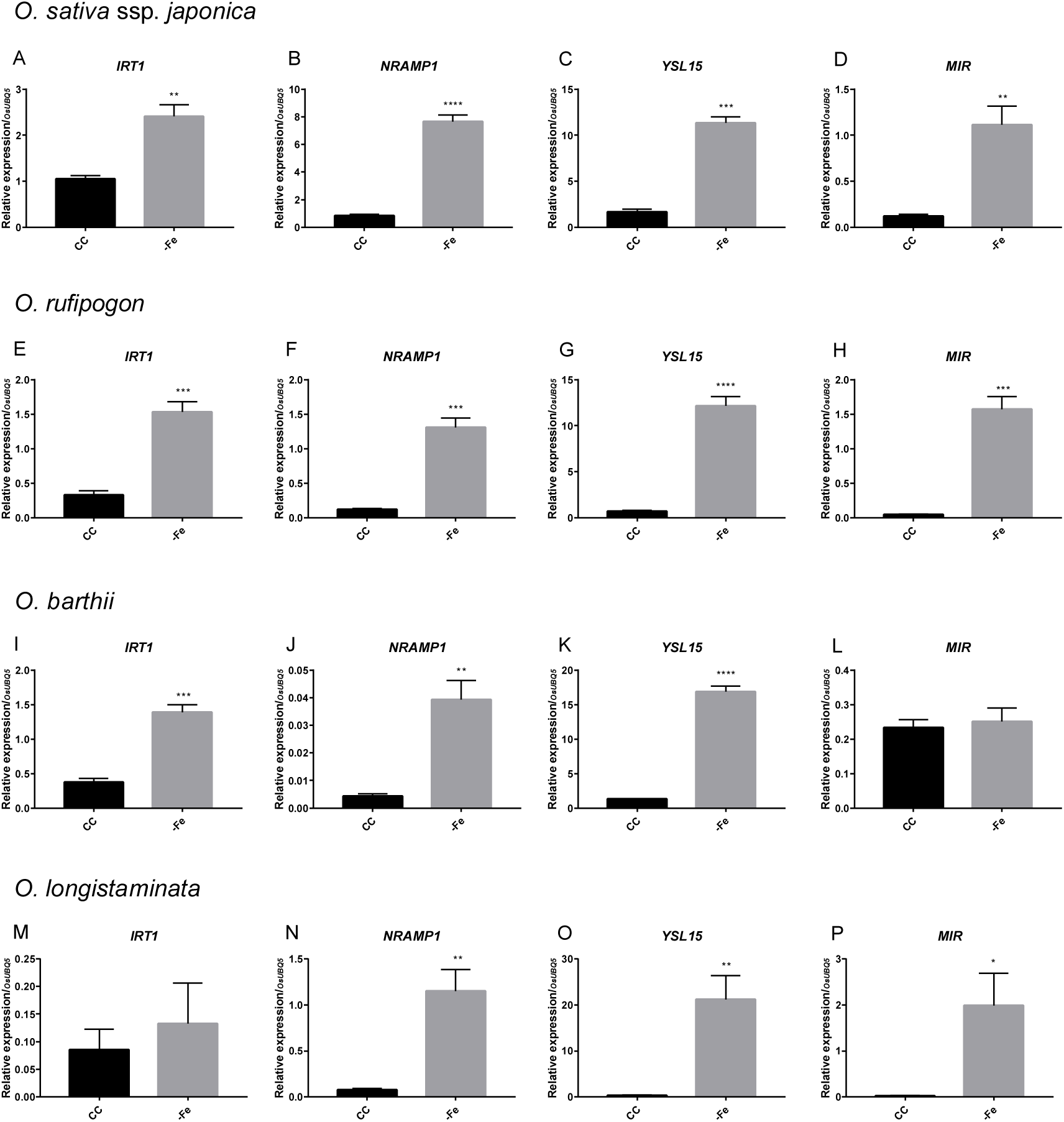
Expression analysis of Fe deficiency-related genes in roots of (A-D) *Oryza sativa*; (E-H) *O. rufipogon*; (I-L) *O. barthii*; (M-P) *O. longistaminata*. Relative expression levels (RT-qPCR, relative to *OsUBQ5* expression) of selected genes (*IRT1*, *NRAMP1, YSL15* and MIR), in roots of plants submitted to control (CC) or Fe deficiency (-Fe) conditions for five days. Roots were collected from plants grown on non-aerated nutrient solution, at three-leaf stage on both conditions at the time of RNA extraction. Values are the averages of four samples (3 plants each) ± SE. Asterisks indicate statistical difference between plants grown under CC and -Fe conditions (Student T-test, *P-value < 0.05, **P-value < 0.01, ***P-value < 0.001, ****P-value < 0.0001).

### Phylogenetic analysis reveals possible origins of *MIR* genes

In order to identify possible distant homologies, we built a DNA hidden Markov model from the identified *MIR* genes – referred from now on as *MIR*-profile – and scanned plant genomes with it. Such methodology is more sensitive than local alignment strategies and hence capable of capturing distant homology relationships among sequences (Pearson 2013). Our approach revealed *MIR*-profile hits in all evaluated grass genomes plus two non-grass monocotyledons: *Musa acuminata* and *Ananas comosus* (Figure 5 and Figure 6). Most of these sequences (≈70%) were found to be part of an exon in genes that belong to the Raffinose Synthase Family, even though there is no detectable sequence similarity between *MIR* and Raffinose Synthase members at the amino acid level. This suggests that DNA indels (insertions or deletions of bases) may be the main source for the observed divergence between MIR and Raffinose Synthase proteins, which seems to have a distant homology relationship without having a related biological function.

**Figure 5.**
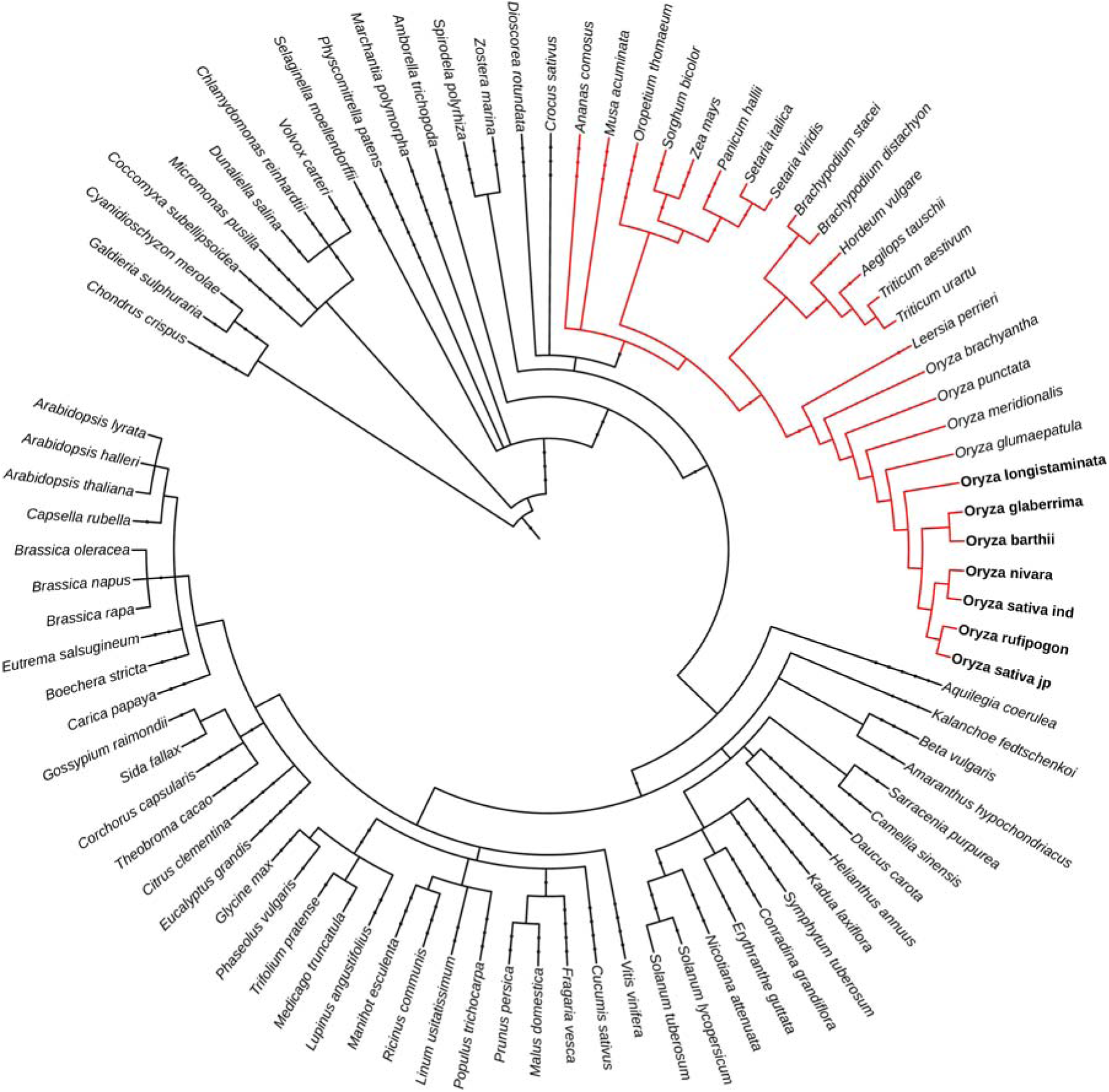
Phylogeny of the plant species genomes screened using the MIR hidden Markov model sequence profile. The red branch comprises species in which a hit was found (E-value < 10^-6^). The species represented in bold are the ones with *MIR* genes. Information on the species phylogeny relationships came from https://www.ncbi.nlm.nih.gov/taxonomy.

**Figure 6.**
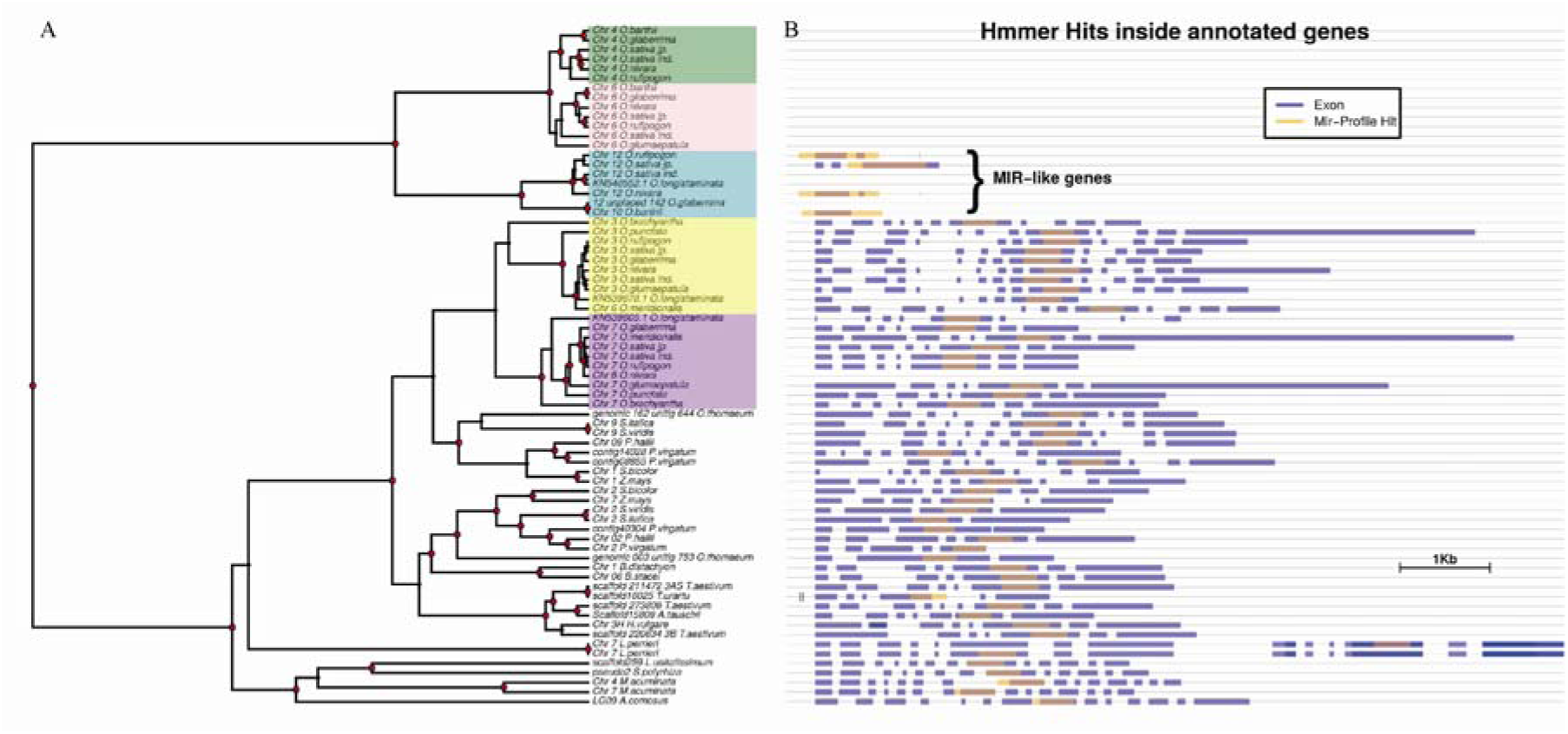
Phylogenetic tree reconstruction of sequences with similarity to MIR and MIR homologs. (A) Reconstructed bayesian phylogenetic tree built using MIR-profile hits. Nodes marked with a red dot indicate support > 0.8. Colored branches show groups of exclusively *Oryza* species. (B) Graphical representation of the position of the MIR-profile hits inside genes (when they occur inside a gene). Exons are shown in blue, MIR-profile hits are shown in orange. Annotated *MIR* genes are highlighted.

We also found highly similar, non-coding sequences in chromosome 4 and chromosome 6 of *O. sativa* ssp. *japonica*, *O. rufipogon*, *O.barthii*, *O. glaberrima* and *O. nivara*. The species *O. glumaepatula*, which does not have a *MIR-like* gene, has a similar sequence only in chromosome 6 (Figure 6B). Based on these sequences plus the Raffinose Synthase hits, we reconstructed a phylogenetic tree (Figure 6B). The tree shows a dichotomy between two major groups. Group I has three subgroups, one containing the *MIR* genes and the other two containing the intergenic segments on chromosomes 4 and 6. These subgroups will be referred respectively as “g-MIR”, “g-Ch4” and “g-Ch6”. Sequences in Group I are derived exclusively from AA *Oryza* genomes: *O. sativa* ssp. *japonica*, *O. sativa* ssp. *indica*, *O. rufipogon*, *O. nivara*, *O. barthii*, *O. glaberrima*, *O. longistaminata* and *O. glumaepatula*. We point out that *O. glumaepatula* is the only species with no sequences in branches g-Ch4 and g-MIR, which may indicate that the genomic fragment similar to *MIR* on chromosome 6 is the one that originated the others by gene duplication events (see Discussion). The second major group, named Group II, clusters *MIR*-profile hits located at an exon of genes that belong to the Raffinose Synthase Family. Among these branches, two of them are exclusively composed by the *Oryza* genus and they will be referred as “g-Ch7” and “g-Ch3”, because most *MIR*-profile hits of each are located, respectively, in chromosomes 7 and 3. Proteins encoded by genes bearing *MIR*-profile hits were scanned against the SMART protein database for domain prediction and all of them were classified as members of the Raffinose Synthase Family (even the ones missing annotation). Annotations for all genes are shown in Figure 6B.

We next confirmed that the genomic blocks in which these sequences similar to *MIR* occur are syntenic to each other inside subgroups g-Ch4 and g-Ch6, but do not share synteny with each other (i.e., g-Ch4 and g-Ch6 sequences for the same species are not in syntenic blocks; Figure 7). We also observed that there is a genomic segment in *O. glumaepatula* chromosome 4 that is syntenic to the genomic blocks in which sequences similar to *MIR* of Ch4 subgroup are present, even though *O. glumaepatula* chromosome 4 itself does not show any sequences with similarity to *MIR*.

**Figure 7.**
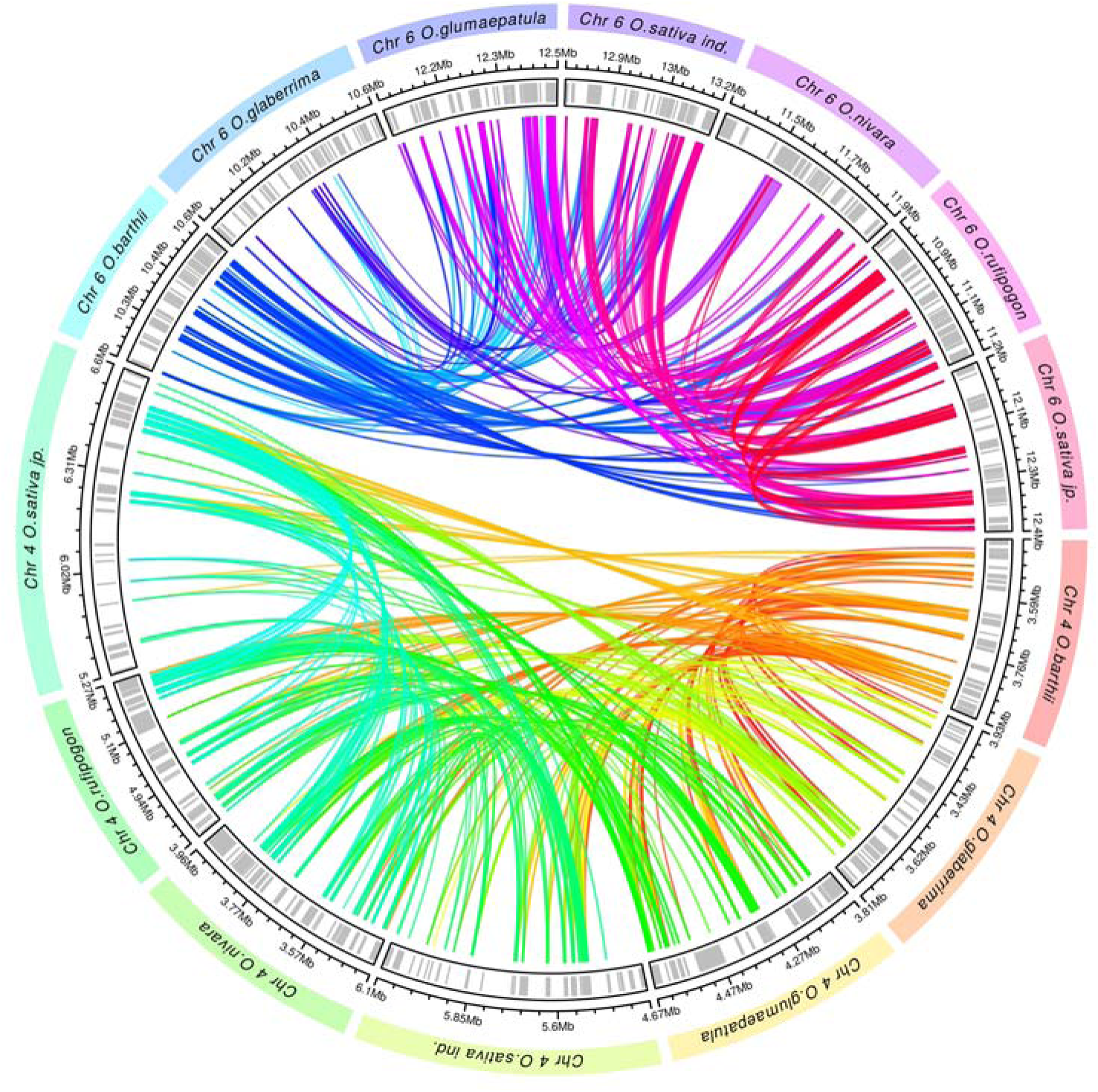
Circular plot of synteny analyses from *MIR-like* sequences in chromosomes 4 and 6 of *Oryza* species. Rectangular arcs forming the circles represent genomic segments. Grey vertical bars inside represent genes. Ribbons connect pairs of genomic loci that are syntenic (E-value < 10E-30).

### The *OsatMIR* co-expression subnetwork cluster is responsive to Fe deficiency

Aiming to shed light on which genes might be functionally associated with *OsatMIR*, we performed a stepwise co-expression meta-analysis applying multiple statistical filters to infer *OsatMIR* subnetwork cluster. Here we used two different microarray platforms (Agilent and Affymetrix) in order to account for oligo DNA probeset bias. By examining the resulting co-expression networks, we can observe that *OsatMIR* lies in the vicinity of genes that are responsive to Fe deficiency regardless of the microarray technology platform employed in building the network (Figure 8A - B). Moreover, Gene Ontology Enrichment analysis of each co-expression networks reveals exclusively biological processes closely related to iron deficiency responses, such as “response to iron starvation” and “iron ion transport”, as well as several methionine related processes, like “methionine biosynthetic process”, “methionine metabolic process” and “L-methionine salvage from methylthioadenosine” (Figure 8C - D). The Agilent network has 239 nodes (genes) and the Affymetrix network has 129 nodes. Overlapping both networks outputs the amount of 38 genes, which therefore can be consistently considered part of the *OsatMIR* co-expression subnetwork (Figure 8G). Notably, 36 out of those 38 genes (plus *OsatMIR*) are induced in plant roots submitted to Fe deficiency treatment according to our differential expression RNAseq analysis (FDR *<* 0.05), which shows that *OsatMIR* is part of a core clustered module responsive to Fe deficiency. In this overlapping cluster we find known genes important for iron homeostasis, such as the transcription factor OsIRO2 (Os01g0952800) (Ogo, et al. 2006), genes related to synthesis and transport of deoxymugineic acid such as deoxymugineic acid synthase (Os03g0237100), IDI2 (Os11g0216900) and IDI4 (Os09g0453800) (Ogo, et al. 2007), the transporter of mugineic acid TOM1/OsZIFL4 (Os11g0134900) (Nozoye, et al. 2011; Ricachenevsky, et al. 2011), genes associated with synthesis and transport of nicotianamine, such nicotianamine synthase (Os03g0307200) and efflux transporter of nicotianamine (Os11g0151500). In addition, the major transporter of Fe^3+^-phytosiderophore complexes, YSL15 (Os02g0650300) was also identified, and the recently described transporter of Fe^2+^-nicotianamine and Fe^3+^-deoxymugineic acid, YSL9 (Os04g0542200) (Senoura, et al. 2017). The list also comprises genes that were already reported by other authors as up regulated under Fe deficiency and down regulated by Fe toxicity, Os01g0871600 and Os01g0871500, both described as TGF-beta receptor, type I/II extracellular region family protein (Bashir, et al. 2014).

**Figure 8.**
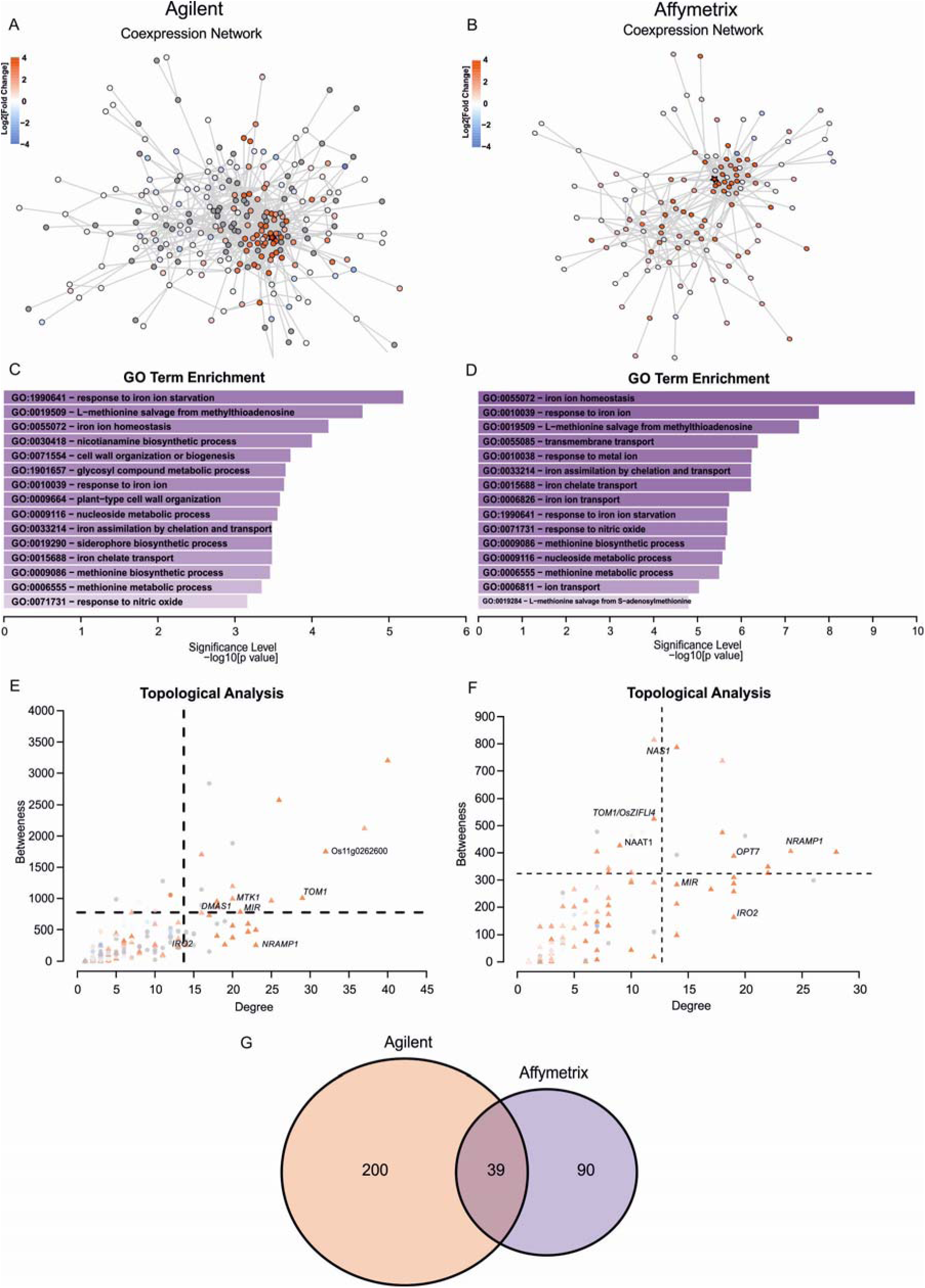
Meta-analysis aiming to elucidate *MIR* co-expression subnetwork. The same steps in the meta-analysis were carried out for two different microarray platform technologies: Agilent (left panel) and Affymetrix (right panel). (A-B): Consensus co-expression networks derived from the co-expression meta-analysis as detailed in Methods. The star-shaped node represents *OsatMIR*. A color gradient (from blue to white to red) was applied on the nodes to denote expression levels in response to Fe deficiency (blue is under-expressed and red is over-expressed). Grey nodes denote probes that were not matched to annotated genes. (C-D): Statistically enriched Gene Ontology Terms on the networks. (E-F): Scatter plot of centrality measurements (degree and betweeness). Coloring scheme is the same as in (A-B). Triangles represent differentially expressed genes in response to iron deficiency and circles are not differentially expressed. Dashed lines are limits for 1 standard deviation above the mean. (G): Venn diagram comparing node composition on both evaluated co-expression networks.

The nodes that share an edge with *OsatMIR* in both Affymetrix and Agilent networks are the ones whose gene expression pattern is tightly correlated with *OsatMIR*’s. Among these genes directly connected to *OsatMIR*, we find Os07g0258400 (*NRAMP1*), Os03g0751100 (*OPT7*), Os03g0307300 (*NICOTIANAMINE SYNTHASE 1*), Os01g0871600 (similar to peptide transporter PTR2-B) and others (Table 2). Furthermore, from a topological point of view, we observed that the group of genes around *OsatMIR* tends to display central positions (either as *Hubs* and/or *Bottlenecks*), which is not surprising, given that the whole meta-analysis was focused on reconstructing the co-expression links around *OsatMIR*. Still, it is worth to highlight that Os07g0258400 (*NRAMP1*), Os03g0751100 (*OPT7*) and Os03g0431600 (hypothetical gene) are highly connected hubs in both Agilent and Affymetrix networks (Figure 8E - F), which suggests that those genes have pivotal roles within the context of this particular subnetwork.

**Table 2.**
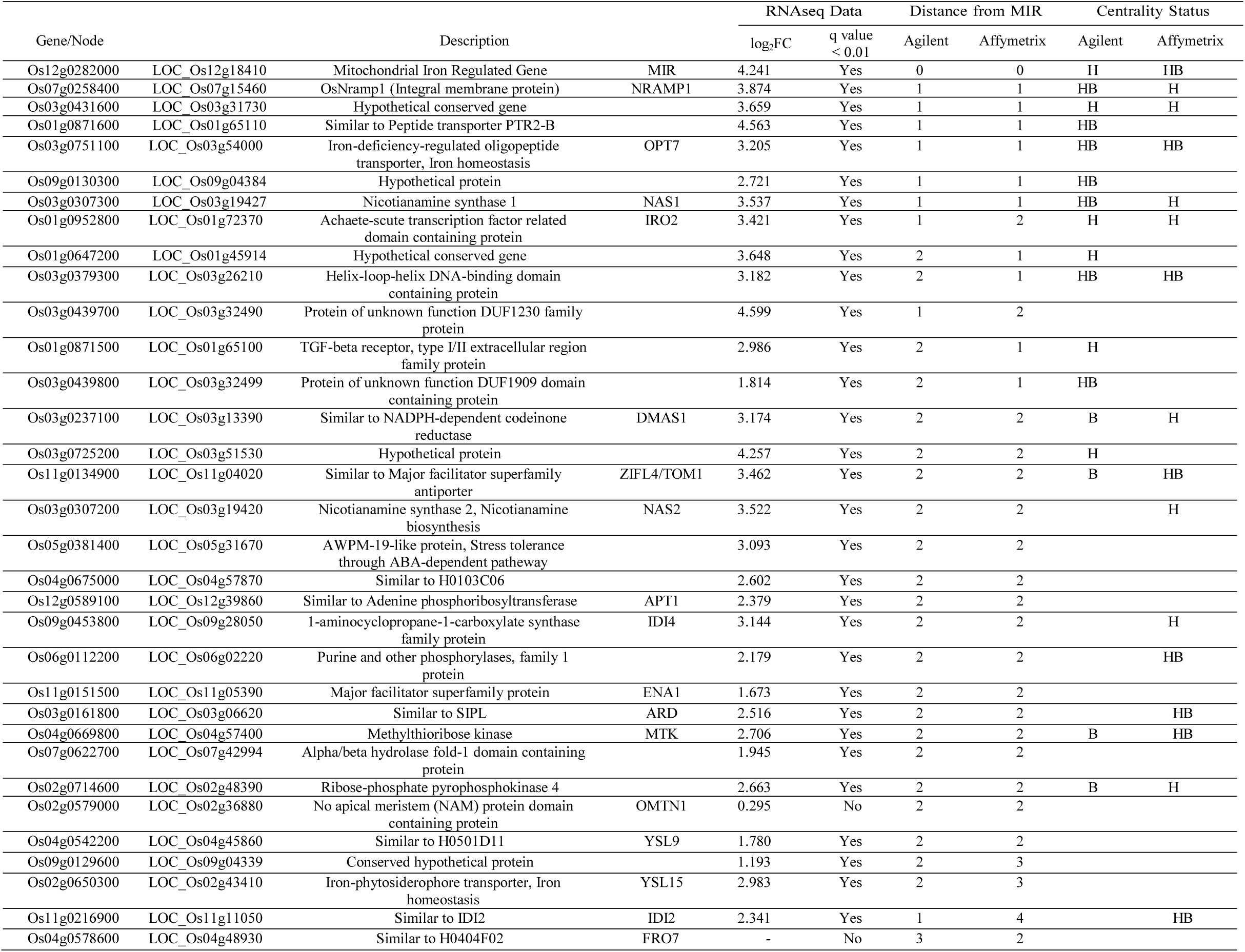
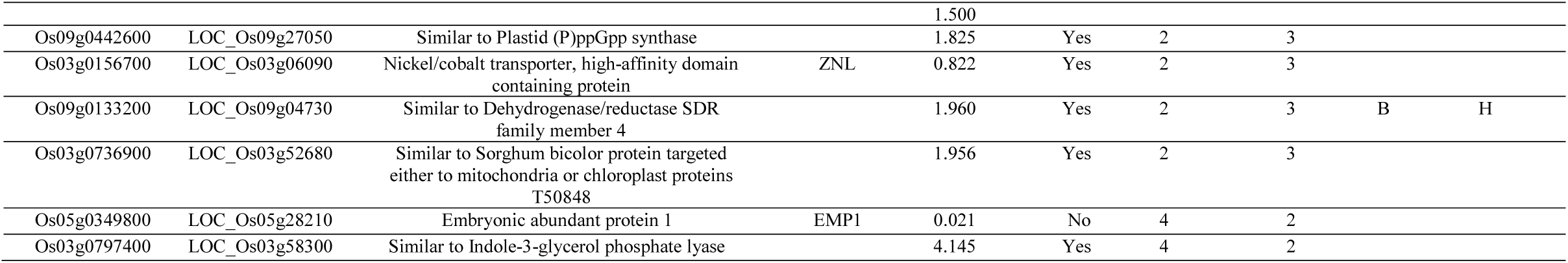
Gene co-expressed with MIR in both microarray platforms.

## Discussion

### *MIR* is not an orphan, but rather a lineage-restricted gene

All sequenced genomes present a set of unique genes. Orphan genes are defined as sharing no similarity with genes or protein domains in other evolutionary lineages (Tautz and Domazet-Loso 2011). The identification of orphan genes depends both on the detection method and the reference set of genomes considered, and thus orphan genes are more likely to be classified as such when only genomes from distantly related species are available. It is hypothesized that, since protein count is fairly constant comparing closely related species, but there is always a certain number of orphan genes, there is equilibrium between gene origin and extinction (Arendsee, et al. 2014). It would also be reasonable to assume that most turnover occurs in young genes (Tautz and Domazet-Loso 2011), which have not yet assumed a central position in metabolism. Thus, genes unique to certain species or lineages can be important for modulating existent pathways, rather than necessary for its function. The *Oryza* genus, which recently had several genomes sequenced (Stein, et al. 2018), is especially attractive for identifying gene families for genes considered orphans based on previous analyses of the rice genome (Zhang, et al. 2019).

Here we showed that, contrary to previous reports (before the availability of *Oryza* wild species genomes), *MIR* is not a rice-specific orphan gene, but part of a new gene family that is taxonomically restricted to a subset of *Oryza* AA genome species. Considering the genomes available, we only found *MIR-like* sequences in AA species, but not in *O. punctata* (BB genome), *O. brachyantha* (FF), *L. perrieri* (commonly used as an outgroup for *Oryza* genus studies) (Stein, et al. 2018), or any other genome available at public databases. The AA species clade within *Oryza* diverged around 2.5 million years ago (MYA), and *MIR* sequences are not present in *O. meridionalis*, a species found only in Australia that diverged later (∼2.4 MYA), or in *O. glumaepatula*, which is found only in South America and diverged later (∼1 MYA); (Stein, et al. 2018). *MIR* sequences were found in both species of the African lineages, *O. barthii* and *O. glaberrima*, in all species of the Asian lineage, and in *O. longistaminata*, which diverged later than *O. glumaepatula* (Reuscher, et al. 2018) (Figure 6A). Since the split between these two lineages occurred right after *O. glumaepatula* speciation, functional *MIR* origination can be timed to less than 1 MYA.

### A model for *MIR* origination

Our data also shows that non-coding sequences similar to *MIR* (which can be detected using BLAST) are found in chromosomes 4 and 6 of several *Oryza* genomes, including *O. glumaepatula*, which lacks functional *MIR* sequences (Figure 6). Chromosomes 4 and 6 sequences are more closely related to each other than to *MIR* genes, indicating that they are products of a duplication event, which likely took place after *O. glumaepatula* speciation, as only the sequence in chromosome 6 is observed in the *O. glumaepatula* genome (Figure 6 and Figure 7). However, we did not find these sequences in chromosomes 4 and 6 of *O. meridionalis* or *O. longistaminata*, a result that can be derived from genome gaps in the draft sequence (Reuscher, et al. 2018; Stein, et al. 2018). Nonetheless, the region where the non-coding sequence similar to *MIR* is found in chromosome 4 of all species is present in the *O. glumaepatula* genome draft sequence, implying that duplication occurred after *O. glumaepatula* split. Whether *O. longistaminata* shares the chromosome 4 and 6 duplication event, and the confirmation that *O. meridionalis* has only the chromosome 6 sequence will need further inspection of a more complete genomic sequence. Although it seems that these non-coding sequences were the source of origin for functional *MIR* genes, it remains to be explored if they have any function in the genome.

Strikingly, we found distant similarities between a *MIR*-based hidden markov model and sequences derived from an exon inside *Raffinose Synthase* Family genes, spread throughout the monocot lineages (Figure 6). These results indicate that *Raffinose Synthase* genes might have been the source of genetic material for *MIR* origin, although in an indirect path: the exons of *Raffinose Synthase* genes generated the sequences present in chromosome 6 (and in chromosome 4 after a duplication event), which in turn were the sources of functional *MIR de novo* origination (Tautz and Domazet-Loso 2011; Zhang, et al. 2019).

If correct, this would imply that the *MIR* evolutionary origin includes a mixture of two proposed models for evolution of orphan (or new, lineage-restricted) genes: (1) duplication-divergence model, in which coding sequences are duplicated and undergo mutations that render one adaptive to a new function, but rather different from the original sequence that BLAST tools would not capture similarities; and (2) *de novo* origination, in which random sequences would combine to form functional sites and would come under regulatory control to produce a transcribed RNA, which would next acquire a functional open reading frame (Tautz and Domazet-Loso 2011). *MIR* evolution would be a result of (1) duplication and divergence from a coding sequence of Raffinose Synthase into a non-coding one in chromosome 6 and (2) duplication of the non-coding sequence, and *de novo* evolution into a protein-coding *MIR* gene. Our model for MIR evolution is summarized in Figure 9.

**Figure 9.**
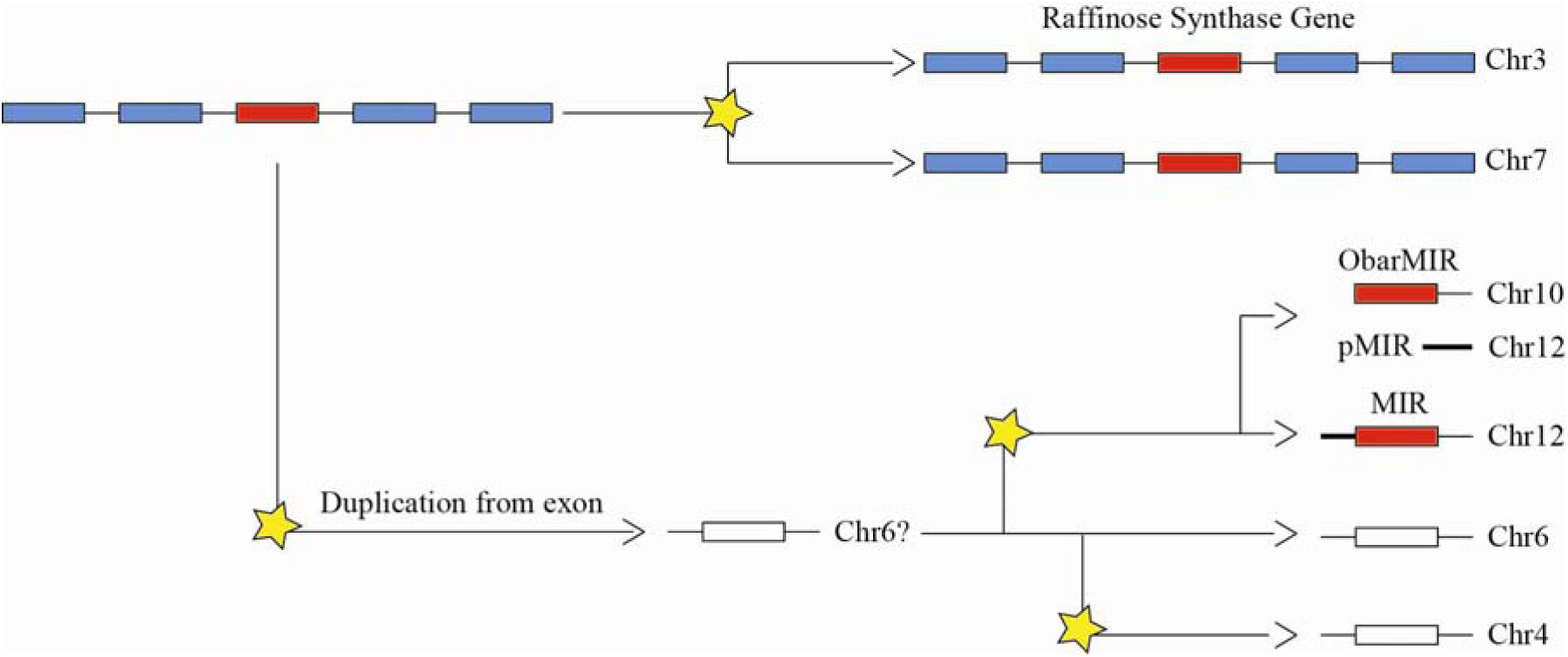
A model for MIR evolution in the monocot lineage. A *Raffinose Synthase* exon was the source of the non-conding sequence present in chromosome 6 of almost all *Oryza* AA genome species (except *O. meridionalis* – which diverged earlier than *O. glumaepatula* - and *O. longistaminata* – which diverged later than *O. glumaepatula*). Given that *O. glumaepatula* has the regions where the non-coding sequence from chromosome 4 and chromosome 6 are found, but has only the chromosome 6 sequence, we suggest that the chromosome 6 was generated first, and was the source of MIR de novo origination - an event that occurred after *O. glumeapatula* speciation. After MIR origination, the sequence in chromosome 6 was also duplicated to generate the non-coding sequence in chromosome 4. The change in chromosomal position of *ObarMIR* and the separation of its regulatory sequence is shown. It should be noted that the Raffinose synthase gene also underwent duplication during the evolution of the monocot lineage. Coding exons are shown as blue boxes; the exon that is related to MIR and a coding sequence are shown as red boxes; non-coding sequences are shown as white boxes; the promoter region of MIR is shown as a thick line. Duplication events are denoted by a yellow star.

### *MIR* function may still be evolving in distinct *Oryza* lineages

*MIR* was previously characterized as functionally important for Fe homeostasis in cultivated rice, and independent expression data has corroborated its role in Fe deficiency responses (Bashir, et al. 2014; Ishimaru, et al. 2009; Stein, et al. 2019; Wairich, et al. 2019) The *MIR* loss-of-function mutant (*mir*) is impaired in correctly perceiving Fe sufficiency status, which leads to increased Fe uptake, up regulation of Fe deficiency response genes, and Fe accumulation (Ishimaru, et al. 2009). MIR protein was localized to mitochondria, and *mir* plants show somewhat similar phenotypic changes to the *mitochondrial iron transporter* (*MIT*) *mit* loss-of-function plants (Bashir, et al. 2011). Based on these studies, *MIR* seems to regulate perception of proper Fe concentrations in mitochondria, which might be important to Fe status sensing. Since Fe sensing is key for making plants more Fe efficient and accumulating Fe in edible tissues for biofortification (Sperotto, et al. 2012), it is important to understand how *MIR* might regulate such function, especially how this protein became entrenched in the Fe regulon, and at which point during *Oryza* speciation *MIR* origination occurred.

We found that most functional *MIR* genes have IDE1-enriched promoters, and are up regulated under Fe deficiency (Figure 1, Figure 4). The exception is *O. barthii*: the *ObarMIR* promoter does not contain IDE1 rich regulatory sequences upstream its coding sequence, and Fe response is impaired (Figure 4). *ObarMIR* is localized in chromosome 10, whereas the regulatory region homologous to *MIR* gene promoters is in *O. barthii*’s chromosome 12, in a region syntenic to *MIR-like* gene localization in all other species (Figure 1 and Figure 2). Thus, the *ObarMIR* coding sequence has changed position within the *O. barthii*’s genome, leaving its promoter in the original location. Interestingly, in *O. glaberrima*, the domesticated species to which *O. barthii* is the progenitor (Wang, et al. 2014), *OglaMIR* also has changed position within chromosome 12, as evidenced by our large-scale genomic alignment (Figure 3). As *OglaMIR* is located in an unplaced scaffold, we were not able to include *O. glaberrima* in MCScanX analyses (Figure 2), due to lack of neighboring genes data - a requirement for MCScan. Thus, we showed that *ObarMIR* was translocated from chromosome 12 to chromosome 10, losing its Fe deficiency-responsiveness due to a lack of IDE1 rich promoter. These results indicate that, although *MIR* might be functional in all *Oryza* species, its role is not central, and can be changed or lost in specific lineages. This corroborates the idea that newly evolved genes might not be central in regulatory networks/metabolism, and therefore might be lost/changed more easily (Tautz and Domazet-Loso 2011; Zhang, et al. 2019). Moreover, based on gene expression data, we have shown that *MIR* genes are likely to have a conserved function in *Oryza* within the subset of AA genome species, where it is found, regulating responses to low Fe.

### MIR co-expression analyses and its relevance to Fe deficiency in *O. sativa*

Our co-expression analysis provided evidence about the importance of *MIR* in response to Fe deficiency and homeostasis in *O. sativa* (Figure 8). Genes described as positive regulators of Fe deficiency response, like *OsIRO2* and the target genes associated with methionine cycle were consistently co-expressed with *OsatMIR.* The induction of methionine cycle under Fe deficiency conditions in grasses is essential for Fe^3+^ uptake by proteins encoded by genes from the *Yellow Stripe* family (Inoue, et al. 2009; Lee, et al. 2009) when complexed with deoxymugineic acid. Furthermore, genes related to long distance Fe transport inside of the body of rice plants, complexed with nicotianamine or not, were also strongly co-expressed with *OsatMIR*, further supporting a central role of *MIR* in Fe homeostasis (Table 2).

Also, among genes that displayed a topological central position on the co-expression networks (meaning they likely play pivotal roles in the *MIR* vicinity subnetwork context), there are three genes that do not present a clear physiological role in Fe deficiency response. These genes were identified as both highly connected hubs and closely associated with *OsatMIR* in both Agilent and Affymetrix networks: *NRAMP1, OPT7* and Os03g0431600 (hypothetical gene) (Table 2). As for *NRAMP1,* studies characterized this gene as Fe^2+^, Mn^2+^, and Cd^2+^ transporter up regulated by Fe deficiency (Takahashi, et al. 2011). In *Arabidopsis thaliana,* the transporter NRAMP1 is important for Fe transport under Fe sufficiency conditions, cooperating with IRT1 to absorb Fe^+2^ from the rhizosphere (Castaings, et al. 2016). The expression level of *iron-deficiency-regulated oligopeptide transporter* (*OPT7*) is specifically up regulated in response to Fe deficiency in root and shoot tissues (Bashir, et al. 2015). Both Fe^3+^-deoxymugineic acid and Fe^2+^-nicotianamine are not substrates transported by OsOPT7. In addition, this gene was not able to complement the *fet3fet4* yeast mutant, which is defective in the Fe uptake when provided with Fe^2+^-nicotianamine (Bashir, et al. 2015; Vasconcelos MW Characterization of the PT Clade of Oligopeptide Transporters in Rice), although it seems to play a significant role in Fe distribution within rice plants (Bashir, et al. 2015). MIR was up regulated in *opt7* lines both in Fe sufficient and Fe deficient medium when compared with wild type plants cultivated under the same condition (Bashir, et al. 2015) corroborating the results obtained by the co-expression network and highlighting the possible concerted role of *MIR* and *OPT7*. Finally, another identified hub closely associated with Osat*MIR* was the hypothetical gene Os03g0431600. According to Ensembl Plant database, this gene does not have an orthologous counterpart in other plant species, however it has three paralogs in the *O. sativa* genome: *Os01g0495701*, *Os06g0294950* and *Os11g0262600*, all of which were up regulated in roots of plants submitted to Fe deficiency for five days (Wairich, et al. 2019). These genes encode proteins ranging from 56 to 59 amino acids long with no predicted conserved domains. To our knowledge, there are no available functional studies for Os03g0431600 and its paralogs and, therefore, they are currently missing meaningful annotation in the major plant databases. However, given previous studies and our co-expression meta-analysis results, we suggest these genes are likely to play an important role in Fe deficiency response and deserve further investigation.

### Conclusion

We propose a model (depicted in Figure 9) in which protein-coding MIR genes are anciently related to a sequence within a *Raffinose Synthase* gene, which by duplication and degeneration resulted in a non-coding sequence with no apparent function. This non-coding sequence was again duplicated and *de novo* evolution resulted in the protein-coding MIR gene around 1 million years ago. *MIR* genes have a function in Fe homeostasis but are still undergoing disruptive evolutionary changes in at least one species (*O. barthii*), including changes in its position and loss of Fe-related regulatory sequences, which can drastically change its role within the plant. Our data provide a detailed example of fast evolution of a newly evolved, lineage-restricted gene, with a functional role in Fe homeostasis in rice and its wild relatives.

## Acknowledgements

This study was financed in part by the Coordenação de Aperfeiçoamento de Pessoal de Nível Superior - Brasil (CAPES) - Finance Code 001, and Conselho Nacional de Desenvolvimento Científico e Tecnológico (CNPq), which granted fellowships to BHNO, AW, JPF and FKR.

